# Same Action, Different Meaning: Neural substrates of Semantic Goal Representation

**DOI:** 10.1101/2021.04.18.440307

**Authors:** Shahar Aberbach, Batel Buaron, Liad Mudrik, Roy Mukamel

## Abstract

Accurate control over everyday goal-directed actions is mediated by sensory-motor predictions of intended consequences and their comparison with actual outcomes. Such online comparisons of the expected and re-afferent, immediate, sensory feedback are conceptualized as internal forward models. Current predictive coding theories describing such models typically address the processing of *immediate* sensory-motor goals, yet voluntary actions are also oriented towards *long-term* conceptual goals and intentions, for which the sensory consequence is sometimes absent or cannot be fully predicted. Thus, the neural mechanisms underlying actions with distal conceptual goals is far from being clear. Specifically, it is still unknown whether sensory-motor circuits also encode information regarding the global meaning of the action, detached from the immediate, movement-related goal. Therefore, using fMRI and behavioral measures, we examined identical actions (either right or left-hand button presses) performed for two different semantic intentions (‘yes’/‘no’ response to questions regarding visual stimuli). Importantly, actions were devoid of differences in the immediate sensory outcome. Our findings revealed voxel patterns differentiating the two semantic goals in the frontoparietal cortex and visual pathways including the Lateral-occipital complex, in both hemispheres. Behavioral results suggest that the results cannot be explained by kinetic differences such as force. To the best of our knowledge, this is the first evidence showing that semantic meaning is embedded in the neural representation of actions independent of immediate sensory outcome and kinetic differences.

**Significance statement:** A standing question in neuroscience concerns the nature of neural circuits representing conceptual information. Previous studies indicate that regions traditionally associated with movement kinematics, also encode symbolic action categories regardless of their specific motor scheme. However, it is currently unclear whether these sensory-motor circuits also play a role in the representation of the intention, for which an action was initiated. Our results demonstrate that an action’s intention, such as its semantic goal, can be discriminated based on neural activity patterns in motor and sensory regions. Moreover, our findings suggest that semantic goals are embedded in sensorimotor regions in a hand-dependent manner.

## Introduction

In everyday life, humans decide, plan and act according to desired goals that span across different time scales and complexities. This remarkable, yet intuitive, trait of structuring goal-oriented actions is at the core of numerous human behavioral control theories. According to predictive coding theories, a corollary discharge induces an embodied simulation of an action’s intended sensory outcome, which is then compared with the actual outcome. This is also known as the *forward model theory*, as it models the causal relationship between actions and their consequences for accurate movement control (Wolpert et al., 1995; Miall and Wolpert, 1996; Wolpert and Flanagan, 2001; Tian and Poeppel, 2010). In line with this theory, previous findings show that during action planning and execution, neural activity in motor regions is continuously modified by the relations between expected and actual sensory outcomes of one’s movement. fMRI findings have demonstrated that similar actions evoke differential neural activity in different regions of the sensory-motor network (including the primary motor, premotor, and parietal cortex) depending on the coupled sensory consequence (Eisenberg et al., 2011; Gallivan et al., 2011; Krasovsky et al., 2014). For example, Krasovsky et al. (2014) show that neural responses evoked by similar horizontal hand movements, depend on the coupled movement direction of a cursor. Additionally, electroencephalography (EEG) recordings have shown that sensory outcomes modulate preparatory motor activity preceding voluntary actions (Reznik et al., 2018).

The forward model theory and corroborating neural evidence have thus far mostly focused on *immediate* sensory-motor goals. For example, generating the correct grip for (immediately) grasping a teapot of a specific load force (Wolpert and Flanagan, 2001). However, voluntary actions are performed to accomplish not only immediate sensory goals, but also (and perhaps even mostly) distal conceptual goals and intentions for which the sensory consequence is sometimes absent or cannot be fully predicted. Moreover, depending on context, similar actions evoking similar immediate sensory feedback can be performed for different (and sometimes even opposite) semantic intentions. For instance, in social interactions, the same actions (e.g. hand gesture, eye-contact and verbal output) can have completely different intended meanings - such as waiving to either say ‘hello’ or ‘goodbye’.

The *grounded cognition hypothesis* addresses the gap between representing immediate and distal goals or intentions by suggesting that central representations in cognition are derived from and depend on modal simulations of actions and perception of sensory consequences (Gallese and Lakoff, 2005; Barsalou, 2008, Kiefer and Pulvermüller, 2012). At the neural level, these simulations are realized as activation patterns in sensory-motor circuits that are similar to those triggered during actual action and perception. Support for this notion comes from studies attributing a cognitive role to motor circuits by showing their involvement in decision making, language (Pulvermüller and Fadiga, 2010; Fernandino et al., 2015; Mollo et al., 2016; Schaller et al., 2017), action selection (Gallivan et al. 2018) and in the semantic representation of objects (Beauchamp and Martin, 2007; Hamilton and Grafton, 2008) and actions (de Lange et al., 2008; Gallivan et al., 2011; Wurm et al., 2016). Nevertheless, to date, the underlying processes that link the physical attributes of actions with their underlying distal intentions are unknown. Therefore, in the current study we set to examine how internal representations of the semantic goals of actions modulate neural activity patterns and behavioral measures. Specifically, using whole brain fMRI, response time and applied force measurements we examined differences between two communicational goals; the intention to respond in affirmation (i.e. responding “yes”) or negation (i.e. responding “no”) using identical actions (either right or left hand button presses). Importantly, the actions were devoid of a difference in immediate sensory outcome.

## Material and Methods

We conducted a behavioral and fMRI study to assess whether movement kinetics and neural activity patterns are modulated by semantic goals in the absence of immediate action-related sensory outcome.

### Participants

Twenty-six subjects (4 males, mean age 23.03, range 18-28 years) participated in the behavioral study, and thirty-three different subjects participated in the fMRI study. Two subjects did not complete the full scanning session due to discomfort in the scanner or difficulty in comprehending the experimenter’s instructions, leaving data from thirty-one participants (16 males, mean age 26.7, range 19-34 years). All participants were healthy, right-handed (self-report), had normal or corrected-to-normal vision, and were naïve to the purposes of the study. The studies conformed to the guidelines approved by the ethical committee in Tel-Aviv University and the Helsinki Committee of the Sheba Medical Center. All participants provided written informed consent to participate and were compensated for their time.

### Experimental design

In the behavioral study, we measured the response time delay and applied force when subjects pressed a button with the intention to affirm (i.e. responding “yes”) or to negate (i.e. responding “no”) a preceding question. In the behavioral study, each trial began with the presentation of a question – either ‘is it a face?’ or ‘is it a vase?’ for 1s, followed by a brief (100ms) presentation of an image. The image was either a silhouette of two profile faces or a central vase (akin to the Rubin vase/face illusion). These images (adapted from the ones used in Hasson et al. (2001) share low-level features (uniform coloring of the object over a striped background), but at short presentations their perception is biased to either a face or a vase. The stimuli were presented at the center of a **23’** monitor, and subtended 3.7° X 4° visual angle. Following image presentation, a question mark appeared on the screen, cueing participants to respond ‘yes’ / ‘no’ at their own pace using either right- or left-hand button-presses according to a predetermined mapping described below. Following participants’ response, a fixation sign was presented at the center of the screen, for a duration completing a 5 seconds Inter Trial Interval (ITI) (see fig.1. for the experimental design).

**Figure.1.**
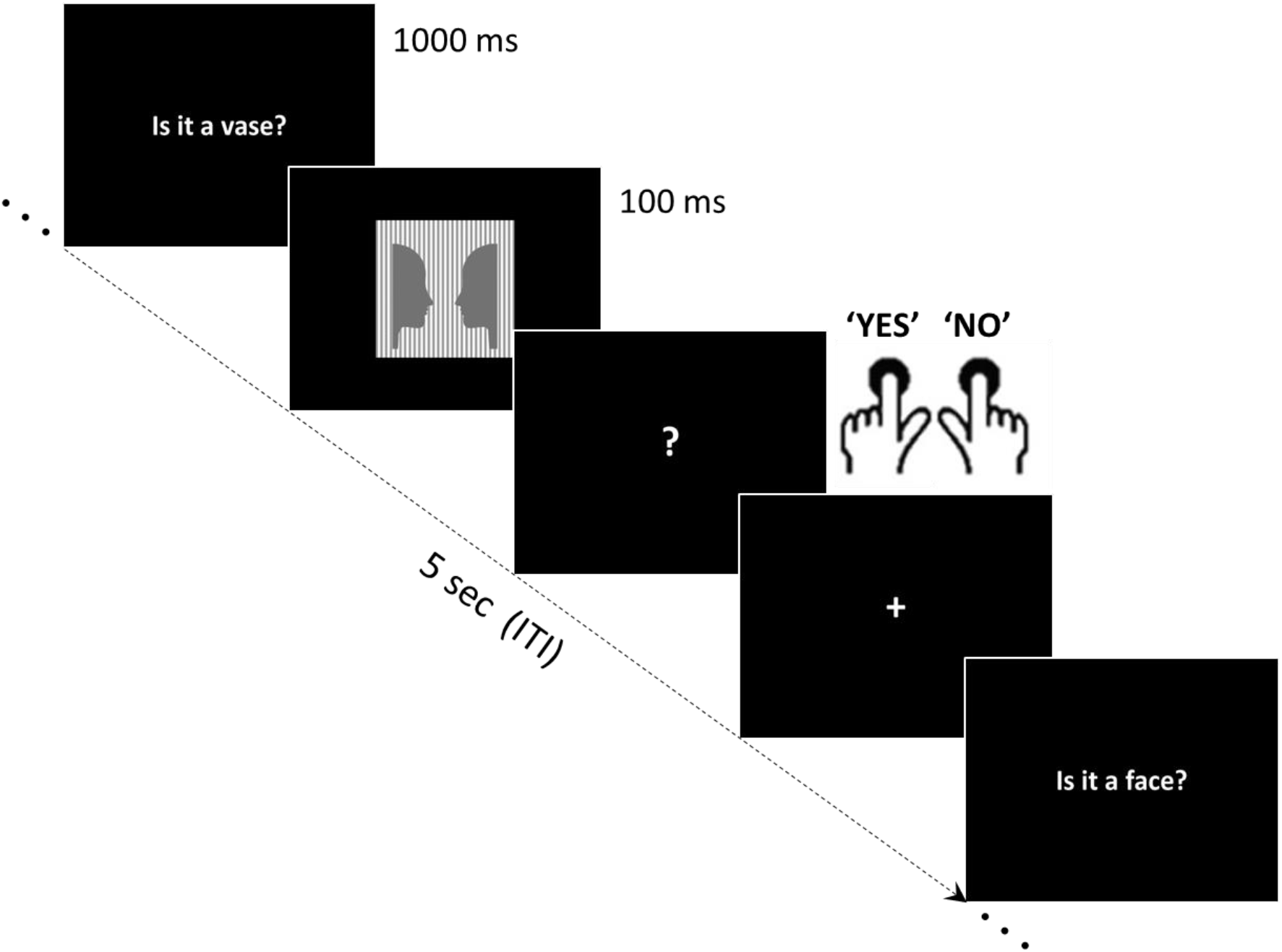
Experimental Trial and Design Example trial of one of the four response type and percept combinations in the behavioral study. This example trial is taken from a right hand =‘no’ / left hand =‘yes’ hand mapping block and requires a ‘NO’ (right hand) response to the question ‘Is it a vase?’. Hand mapping is fixed once at the beginning of each behavioral block. The sequence of events is similar for the fMRI experiment (for exact time durations see Methods).

The experiment included four experimental runs of 80 trials each. In two runs, the right-and left-hand index fingers were mapped to ‘yes’ and ‘no’ responses respectively, while the mapping was reversed in the other two runs. Run order was counterbalanced across participants. The different images and question types were counterbalanced such that, overall, within a particular hand there was no consistent difference in sensory input between responses of different semantic meaning. To ensure that participants correctly recalled the mapping between hand (right/left) and response (‘yes’/’no’), they performed 20 practice trials before each run.

The experiments were performed using Psychtoolbox-3 (www.psychtoolbox.org) on MATLAB 2016b (The MathWorks, Inc., Natick, Massachusetts, United States). Applied force was measured using force sensors (FlexiForce™ A301, Tekscan Inc., Boston, MA) with a dynamic range up to 4.4N and repeatability of constant force measurements of <0.025 N. The sensors were placed under two rubber buttons and connected to analog pins on Arduino^®^ mega2560. The signal from each sensor was read using MATLAB Support Package for Arduino Hardware at a rate of 25Hz. Press onset and offset were detected using a threshold of 0.28N.

The experimental design of the fMRI study was similar to the behavioral study described above, with a few modifications. We used a longer ITI (12s) and a shorter time for question presentation (900ms). Subjects were required to respond within a time window of 2s in order to maintain time-locking between subsequent stimulus presentation (size 1.13°X 1.24°) and scanner acquisition time. No force measures were obtained in the scanner. Prior to each run, subjects performed 16 practice trials with the corresponding hand mapping. Each experimental run contained 48 trials.

### fMRI Data Acquisition

Functional imaging was performed on a Siemens Magnetom Prisma 3T Scanner (Siemens Healthcare) with a 64-channel head coil at the Tel-Aviv University Strauss Center for Computational Neuroimaging. In all functional scans, an interleaved multiband gradient-echo echo-planar pulse sequence was used. Whole-brain coverage was provided by acquiring 66 slices for each volume (slice thickness 2 mm; voxel size 2 mm isotropic; TR= 1000ms; TE=30ms; flip angle = 82°; field of view = 192mm; acceleration factor = 2). For anatomical reference, a whole-brain high-resolution T1-weighted scan (slice thickness 1mm; voxel size 1mm isotropic; TR= 2530ms; TE= 2.99ms; flip angle= 7°; field of view = 224mm) was acquired for each participant.

### Statistical Analysis

#### Behavioral session

Differences in delay between cue (appearance of question mark) and participants’ response (response time; RT), differences in button-press peak force and in the force area under the curve (AUC) across conditions were compared using a three-way repeated measures ANOVA with hand (right/left), response type (‘yes’/’no’) and percept (face/vase) as within-subjects variables. Behavioral data were analyzed using SPSS Statistics version 27 (IBM). and *JASP* (JASP Team 2020, Version 0.14.1) was used for Bayesian analysis when required.

#### fMRI session

fMRI data preprocessing and first-level GLM analysis were conducted using the FMRIB’s Software Library’s (FSL v5.0.9) fMRI Expert Analysis Tool (FEAT v6.00) (Smith et al., 2004). The data from each experimental run underwent the following pre-processing procedures: brain extraction, slice-time correction, high-pass filtering at 100s (0.01 Hz), motion-correction to the middle time-point of each run, and correction for autocorrelation using pre-whitening (as implemented in FSL). Trials with head motion that exceeded 2 mm were excluded from further analysis (max 8 trials within a subject). All images were registered to the high-resolution anatomical data using boundary-based reconstruction and normalized to the Montreal Neurological Institute (MNI) template using nonlinear registration. Anatomical regions were identified using the Harvard-Oxford cortical structural atlas and the Automated Anatomical Labeling (AAL) atlas.

In order to detect differences in spatial patterns of activity across conditions, we used a multivariate testing approach (Multi-t) (Gilron et al., 2017) which has been shown to have better statistical power relative to decoding methods (Rosenblatt et al., 2019). We used the Multi-T analysis with a whole-brain searchlight approach, similar to the one employed in Krasovsky et al. (2014), to discriminate activation patterns across conditions (e.g., yes/no responses within each hand). For each voxel and each trial, we calculated activity level as the percent signal change relative to the time course mean. To take into account the hemodynamic delay, we used the 5th TR from the button-press onset. Compatible with the number of trials, we obtained for each voxel 192 values across all experimental conditions (48 for each combination of hand and semantic goal). The number of trials within conditions varied slightly across participants due to response errors in the task (slow responses or using the wrong hand) or excessive movement in the scanner. In order to keep the trial number identical across conditions, for statistical comparisons within each participant, we randomly sampled N trials from conditions with more trials, where N represents the number of trials in the condition with least number of trials for that subject.

For each voxel, defined as a center-voxel we outlined a neighborhood which included the center voxel and its 26 closest voxels using Euclidean distance. The activity patterns in this neighborhood from all trials were compared across conditions and the central-voxel was assigned with a corresponding multivariate-t value. In addition, for each participant, we shuffled the data labels according to the relevant test (e.g. right/left hand) and repeated the same analysis that was performed on the data using the original labels. Overall, for each participant, we obtained a map of real data t-values and 400 maps of t-values based on the shuffled data.

To determine group-level significance, we used the permutation scheme suggested by Stelzer et al. (2013). First, we averaged all the real statistical maps across subjects to create a group average map. Next, we randomly chose one shuffle map from each participant and averaged those shuffled maps across participants to create one average shuffled map. We repeated the procedure with shuffled labels 10,000 times, providing a distribution of shuffled data accuracy maps (representing the null hypothesis). Within each voxel we used the distribution of shuffled map t-values to compute a corresponding voxel-wise p-value of the t-value obtained in the real map (lowest possible p-value 1/10,000). We then submitted these p-values to false discovery rate (FDR) correction (Benjamini and Hochberg, 1995), with q = 0.05, to create a binary map of significant voxels.

## Results

### Behavioral study

All behavioral measures were analyzed using a three-way repeated measures ANOVA with hand (right/left), semantic goal (‘yes’/’no’) and percept (face/vase) as within-subjects variables. Significant interactions were further probed using post hoc pairwise comparisons, and corrected for multiple comparisons using Bonferroni correction.

Main effects were found in response time (RT) for all three levels: hand, semantic goal and percept, such that right hand responses were shorter compared to the left (0.86±0.03s vs. 0.9±0.03s; *F*(1,25) = 6.87, *p* =0 .015), ‘yes’ RTs were shorter than ‘no’ (0.81±0.02s vs. 0.94±0.03s; *F*(1,25) = 82.45, *p* < 10^−3^) and responses to a face percept were shorter compared to a vase (0.86±0.02s vs. 0.9±0.03s; (*F*(1,25) = 21.9, *p* < 10^−3^). In addition, an interaction was found between hand identity and semantic goal (*F*(1,25) = 9.35, *p* = .005) such that within the right hand, ‘yes’ RTs were shorter than ‘no’ (0.75±.03s vs. 0.97±0.03s; *t*(25) = 7.67, *p* < 10^−3^, Bonferroni corrected), while in the left hand, no significant difference was found for semantic goal (‘yes’ = 0.87±0.03s; ‘no’ = 0.92±0.03s; *t*(25) = 1.635, *p* =.115). An additional interaction was found between semantic goal and percept (*F*(1,25) = 7.02; *p* = .014) such that within the ‘yes’ semantic goal, RTs for face were shorter than for vase (0.77±0.03s vs. 0.85±0.03s; t(25) = 3.83, p = .001, Bonferroni corrected), while no significant difference was found for percept within the ‘no’ semantic goal (face = 0.94±0.03s; vase = 0.95±0.03; *t*(25) = 1.03, *p* =.312). No interaction was found between hand identity and percept, nor an interaction of hand identity X semantic goal X percept (see Table 1 for full statistical results).

**Table.1.** Statistical results – behavioral experiment Summary of the three-way repeated measures ANOVA results for each of the three behavioral measures: RT (response time), force peak and AUC (area under the curve); behavioral study.

| Predictor | | | $f(1,25)$ | $p$ | $\eta p^2$ |
| --- | --- | --- | --- | --- | --- |
| RT | Hand | Right < Left | 6.87 | .015 | .22 |
|  | Goal | 'yes' < 'no' | 82.45 | .000 | .77 |
|  | Percept | 'face' < 'vase' | 21.90 | .000 | .47 |
|  | Hand * Goal |  | 9.35 | .005 | .27 |
|  | Hand * Percept |  | .20 | .655 | .01 |
|  | Goal * Percept |  | 7.02 | .014 | .22 |
|  | Hand * Goal * Percept |  | .00 | .999 | .00 |
| Force<br>Peak | Hand | Right > Left | 91.94 | .000 | .79 |
|  | Goal |  | .31 | .584 | .01 |
|  | Percept |  | .02 | .884 | .00 |
|  | Hand * Goal |  | .01 | .925 | .00 |
|  | Hand * Percept |  | 1.59 | .219 | .06 |
|  | Goal * Percept |  | .10 | .753 | .00 |
|  | Hand * Goal * Percept |  | .87 | .361 | .03 |
| Force<br>AUC | Hand | Right > Left | 17.36 | .000 | .41 |
|  | Goal |  | .05 | .827 | .00 |
|  | Percept |  | .07 | .797 | .00 |
|  | Hand * Goal |  | .77 | .388 | .03 |
|  | Hand * Percept |  | 1.77 | .195 | .07 |
|  | Goal * Percept |  | .17 | .687 | .01 |
|  | Hand * Goal * Percept |  | 1.42 | .245 | .05 |

For force measures, a significant effect was found for hand identity such that right-hand force peak and AUC were greater compared to the left-hand (Peak: 1.78±0.13N vs. 1.28±0.14N; *F*(1,25) = 91.9, *p* = .000, AUC: 0.58±0.06N/s vs. 0.45±0.07N/s; *F*(1,25) = 17.36, *p* = .000 see Fig.2). No other significant main effects or interactions were found in force measures (see Table 1&2 for a complete description of the behavioral results). Importantly, results from Bayesian paired sample t-test analysis show that for both hand presses, participants exert similar force for the two semantic action goals. This was found both in force peak (Right hand BF_01_ = 4.15, Left hand BF_01_ = 3.62) and AUC (Right hand BF_01_ = 4.69, Left hand BF_01_ = 4.46). The results suggest that semantic action goals do not modulate force measures of the executed action.

**Figure.2.**
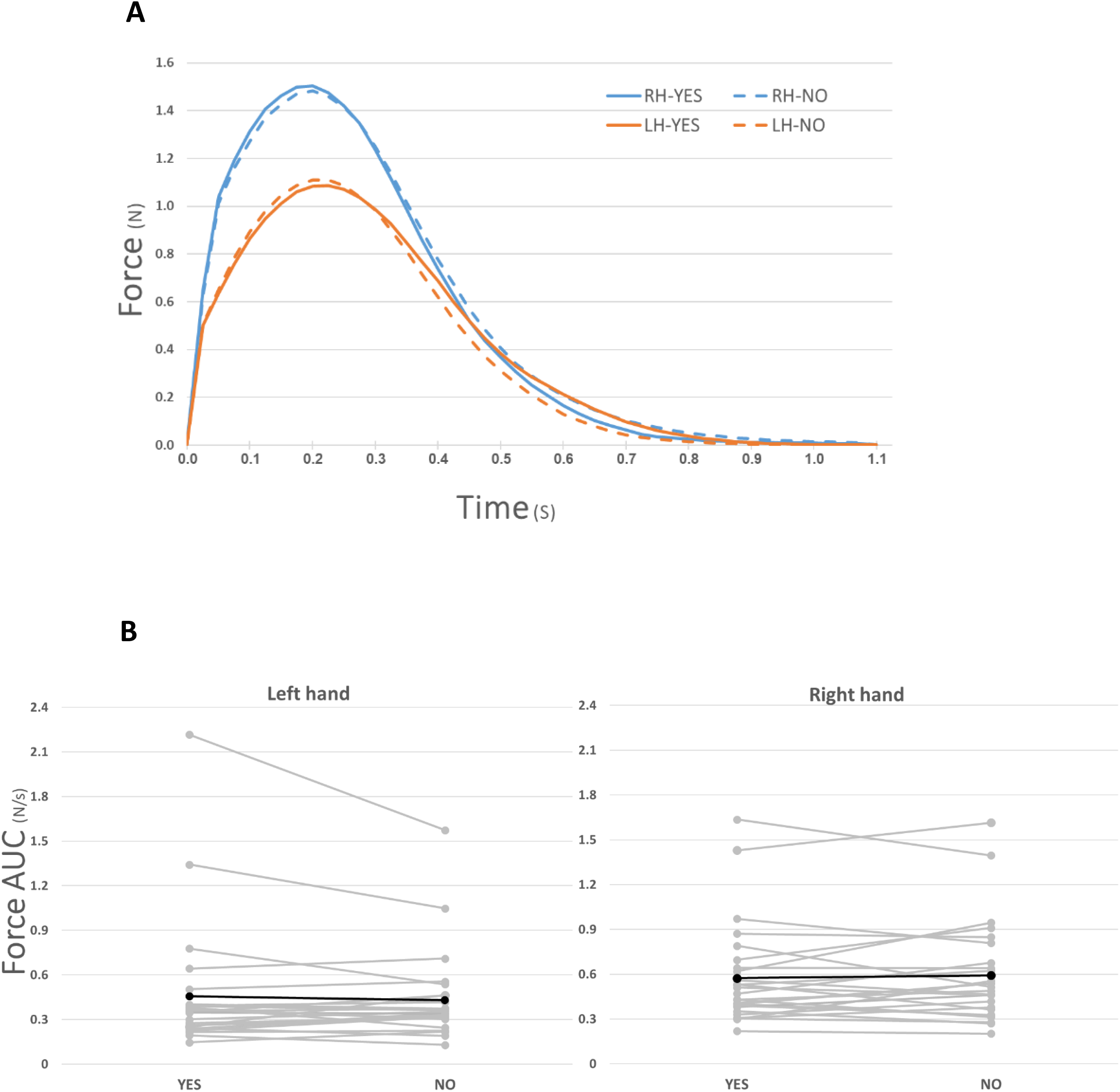
Applied force for semantic goals in the right and left hand Group average force profiles (A), and individual subject’s AUC (B) of ‘yes’ vs. ‘no’ responses within each hand, collapsed over sensory input. Greater force was applied in right hand responses compared to left (**p<0.001), while no difference was found in force levels between the two semantic goals within each hand. RH/LH correspond to Right hand and Left hand respectively.

**Table.2.** Descriptive measures – behavioral experiment: Estimates (mean, with SE in parentheses) of all experimental conditions for the three behavioral measures: RT (response time), force peak and AUC (area under the curve); behavioral study.

|  |  | **RT (s)** |  | **Force peak (N)** |  | **Force AUC (N/s)** |  |
| --- | --- | --- | --- | --- | --- | --- | --- |
|  |  | 'yes' | 'no' | 'yes' | 'no' | 'yes' | 'no' |
| **Right hand** | Face | .71(.03) | .97(.03) | 1.77(.14) | 1.77(.13) | .56(.07) | .59(.06) |
|  | Vase | .79(.03) | .97(.04) | 1.81(.14) | 1.77(.13) | .58(.06) | .59(.06) |
| **Left hand** | Face | .83(.03) | .92(.03) | 1.31(.16) | 1.27(.12) | .46(.08) | .43(.05) |
|  | Vase | .92(.04) | .93(.03) | 1.28(.16) | 1.27(.13) | .45(.08) | .43(.06) |

### fMRI study

In order to detect center-voxels sensitive to the responding hand (right/left), we performed our MVPA multi-T analysis contrasting right- and left-hand trials (collapsed across semantic goals). As expected, results from this analysis revealed significant voxels in the motor strip in the precentral and postcentral gyrus (light blue voxels in figure 3). In order to test for sensitivity for semantic goals, we compared activity patterns evoked by button presses representing ‘yes’ vs. ‘no’ answers separately within each hand. The number of trials within conditions varied slightly across participants (range: 34 - 48 trials) due to trial rejection according to the aforementioned rejection criteria (see methods). Within right hand responses for the two semantic goals (i.e. responding ‘yes’ vs. ‘no’) we found significantly different activity patterns in the right inferior frontal gyrus, the premotor cortex in bilateral precentral gyrus (preCG), left superior parietal lobule (SPL), left angular gyrus, bilateral fusiform gyrus and in the inferior lateral occipital cortex (LOC) bilaterally (*p* < 0.05 FDR corrected, see Figure 3 and Table 3 for coordinates). For the left hand trials, a similar analysis of pattern separation based on semantic goal did not yield voxels that survived correction for multiple comparisons. However, using a more liberal threshold (*p* < 0.0001 uncorrected), revealed voxels in locations adjacent to the ones found for the right hand, in the left premotor cortex and left SPL and voxels that overlapped with those found for right hand trials in the inferior LOC bilaterally (see Figure 4).

**Figure.3.**
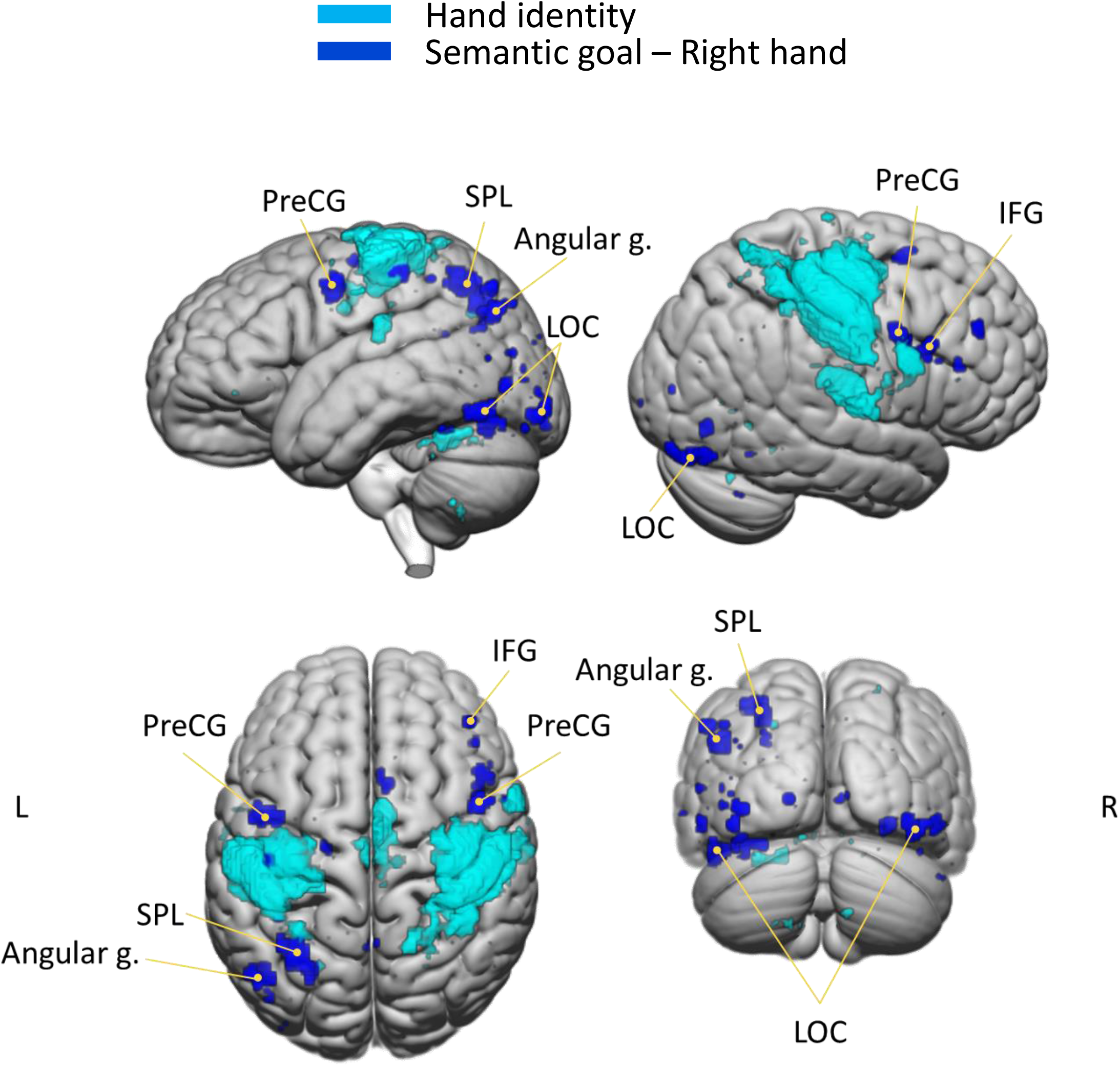
Areas sensitive to hand identity and semantic goals Areas sensitive to hand identity (light blue) in primary motor and supplementary motor area - precentral and postcentral gyrus, bilaterally. Areas sensitive to semantic goals in right hand trials (dark blue) in premotor cortex - precentral and middle frontal gyrus bilaterally left angular gyrus and SPL, and in the LOC and fusiform gyrus bilaterally. p < 0.05 FDR corrected.

**Table.3.**
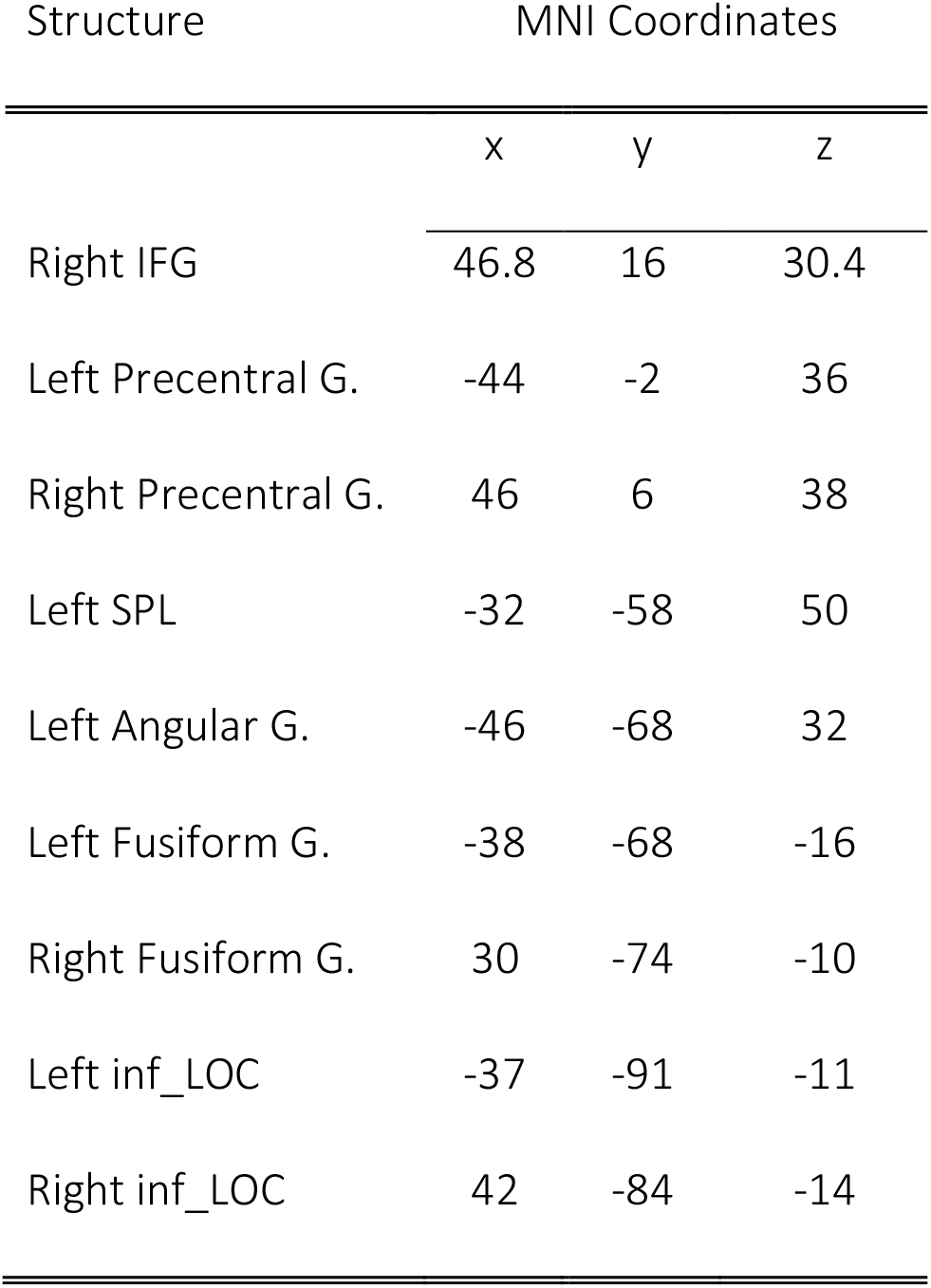
Center coordinates of regions discriminating ‘yes’ vs. ‘no’ semantic goals Center positions (in MNI coordinates) of significant areas sensitive to semantic goals in right hand trials (p < 0.05 FDR corrected).

| Structure | MNI Coordinates |  |  |
| --- | --- | --- | --- |
|  | x | y | z |
| Right IFG | 46.8 | 16 | 30.4 |
| Left Precentral G. | -44 | -2 | 36 |
| Right Precentral G. | 46 | 6 | 38 |
| Left SPL | -32 | -58 | 50 |
| Left Angular G. | -46 | -68 | 32 |
| Left Fusiform G. | -38 | -68 | -16 |
| Right Fusiform G. | 30 | -74 | -10 |
| Left inf_LOC | -37 | -91 | -11 |
| Right inf_LOC | 42 | -84 | -14 |

**Figure.4.**
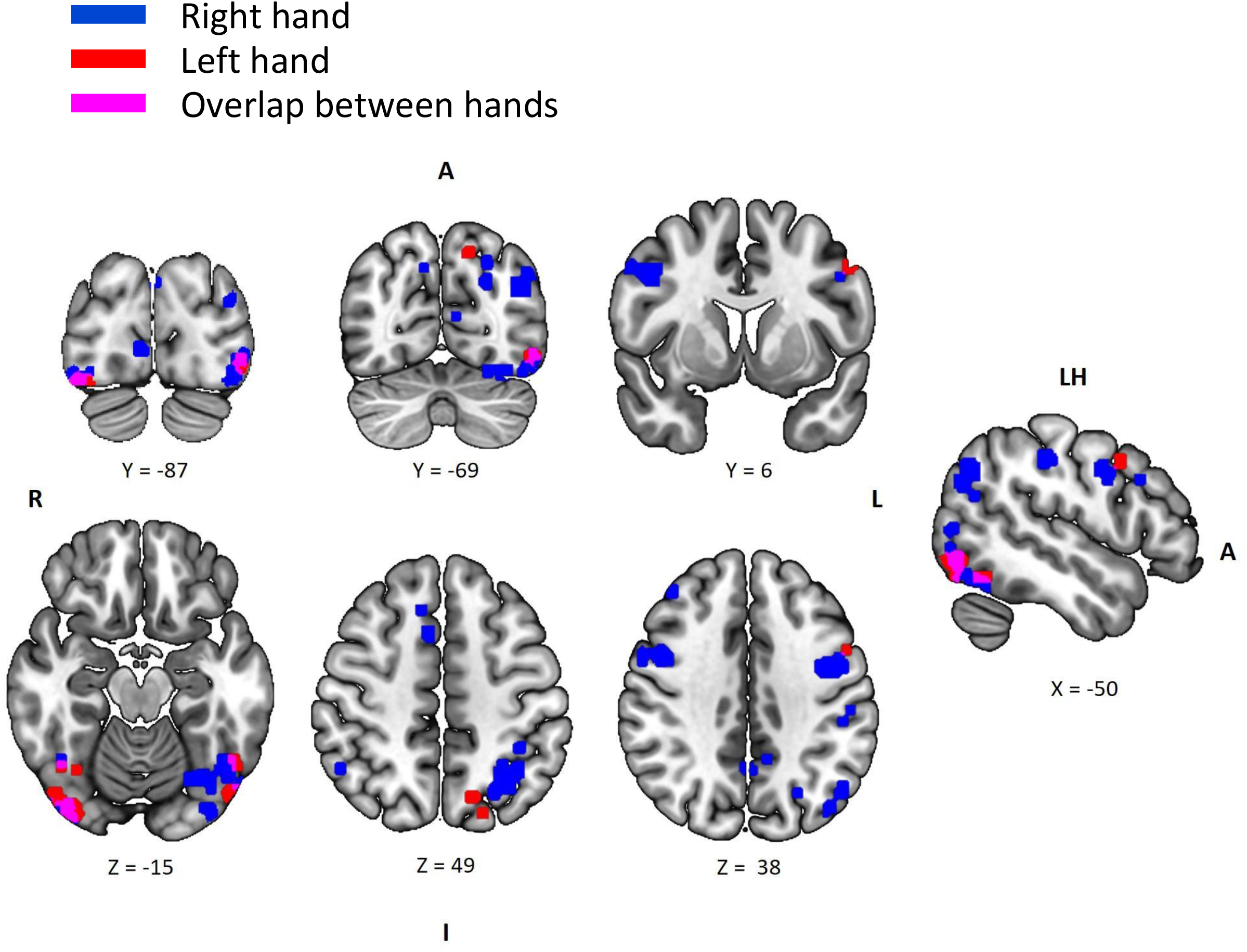
Areas sensitive to semantic goals in both hands Adopting a liberal threshold, areas sensitive to semantic goals in left hand trials (red, p < .0001 uncorrected) are adjacent to significant areas sensitive to semantic goals in right hand trials (blue, p < 0.05 FDR corrected., see fig.3) in left premotor cortex, left SPL, and overlap in the inferior LOC bilaterally (magenta).

In principle, the differences we find in neural activity patterns for semantic goals (yes/no) could be ascribed to differences in RT (Yarkoni et al. 2009). Indeed, similar to the results obtained in the behavioral study, RTs in the scanner during ‘yes’ responses with the right hand (M = .734s, SD = .026) were shorter than ‘no’ responses (*M* = .945s, *SD* = .045; *t*(30) = 7.74, p < 10^−3^, Bonferroni corrected), while no differences were found in the left hand (‘yes’, M = .853s, SD = .043; ‘no’, M = .846s, SD = .03; t(30) =.375, *p* = .710). Therefore, to rule out this potential alternative explanation, the neural activity patterns were also tested for differences due to response times. For each participant, within each hand, trials across all runs were categorically separated to fast/slow according to RT speed using median split, collapsed over semantic goal (i.e. ‘yes’/no’). Importantly, this analysis did not reveal significant voxels of separation, nor an overlap with the areas found sensitive to semantic goals for right hand trials, even when using a more liberal threshold of p = 10^−4^, uncorrected. Another potential explanation for the neural differences we find for yes/no responses could be differences in the force applied during button press. Although in the scanner we did not measure press force, this account is less likely since in the behavioral study we found no significant difference in press-force for yes/no responses (see Figure 2). Therefore, taken together, the behavioral and imaging results suggest top-down, goal-dependent modulation of brain activity both in visual and motor regions despite similar action kinetics and immediate sensory consequences.

Finally, with respect to the neural representation of semantic goals, we examined whether it is hand dependent or rather, can be generalized across the right and left hands. To this end, we used a support vector machine (SVM) classifier (Chang & Lin, 2011) with a two-fold cross validation. The classifier was trained to discriminate activation patterns from yes/no trials based on data from right hand trials and tested on yes/no activation patterns from left hand trials. This analysis was conducted on all significant right-hand voxels that were found in the multi-T analysis described above. However, the resulting accuracy values (voxels’ accuracy range: 0.457 - 0.536) were not significantly higher than the baseline distribution generated by the same analysis performed on shuffled data labels (voxels’ accuracy range: 0.497 - 0.502).

## Discussion

In the current study, we examined how neural activity pattern represent different semantic goals of identical actions. Our fMRI results show that different semantic intentions modulate the neural activity patterns of identical actions in sensory-motor areas in frontoparietal and occipital cortex. Specifically, for right hand trials, a significant separation of ‘yes’ vs. ‘no’ responses was found in the bilateral premotor cortex, left angular gyrus, left SPL, inferior occipital cortex (LOC) and fusiform gyrus bilaterally. Similar, although weaker, patterns were found for the left hand. Importantly, the neural patterns found in the current study are independent of immediate sensory consequence as there was no consistent difference between ‘yes’ and ‘no’ button presses with respect to the sensory feedback. Moreover, the differential activity patterns found for the two semantic goals are independent of kinetic differences as no difference was found in pressing force between the execution of the two goal-directed movements when tested in the behavioral study.

Neural activity in the premotor cortex and intraparietal regions were previously shown to encode a variety of action-related goals (Hamilton and Grafton, 2006; Moritz et al., 2015; Gallivan et al., 2016; Gertz et al., 2017). For example, the performed grip type (a whole-hand or precision grip, Turella et al., 2020) and the target location of the movement (Gallivan et al., 2011) were successfully discriminated based on activity patterns elicited in these areas. Here we show that similar frontoparietal regions encode action-related semantic goals, beyond their involvement in the sensory representation of the visual target or the motor representation of the movement goal. Additionally, high order visual areas including the fusiform gyrus have been shown to selectively represent body parts not only when presented visually but also during action execution with no action-related visual input (Orlov et al. 2010). Here we show that beyond information regarding how or what action was performed, lateral occipital cortex also encode the action’s *intention* for which a movement was initiated.

Compatible with previous findings (Wentura, 2000; Brouillet et al., 2010), our results from response times in the behavioral and fMRI studies show that within the right hand, responses were faster for ‘yes’ compared to ‘no’ answers. It was previously highlighted that the fMRI blood oxygenation level dependent (BOLD) signal is correlated with reaction time latencies in several grey and white matter regions of the brain (Yarkoni et al. 2009). Although the current experimental design did not employ a speeded reaction task, and subjects responded at their own pace, they were still constrained by a 2s time window to respond (in order remain in sync with experimental timing). Therefore, the current imaging results might be affected by differences in response latencies. However, when tested directly by comparing the neural activity patterns of short and long RTs (based on median split of response latencies), no significant voxels were found. Importantly, there was no overlap in voxels sensitive to response time and semantic goal separation within the right hand trials even at a more liberal threshold. Thus, it is less likely that the differences found in the neural activity for semantic goals can be accounted for by differences in reaction time.

The significant separation of activity patterns for the different semantic goals was found for right hand button presses. An exploratory analysis, using a more liberal threshold for the separation within left hand button presses, descriptively show that the areas with highest sensitivity for semantic goals are adjacent to or overlap the areas found most sensitive to semantic goals for right hand button presses. Since the amplitude of the BOLD signal in sensory-motor areas is positively correlated with the level of force used (Sulzer et al., 2011), the failure of left-hand trials to reach significant separation in the multi-T analysis might be due to lower signal to noise ratio (SNR) of left-hand responses. Indeed, it was found in our behavioral study that the pressing force exerted by the subjects was significantly lower in the left compared to the right hand.

The spatial similarity of voxels differentiating semantic goals in right- and left-hand trials, suggests that the two hands might share a common infrastructure representing semantic goals. To address this question, we performed cross-classification within the voxels that were found sensitive to the semantic goal of right hand button presses. We trained a model on right hand trials (‘yes’ vs. ‘no’) and examined classification of left hand trials. Despite the spatial similarity of voxels between hands, our analysis did not yield significant cross-classification. This could imply that these regions differentiate the intended meaning of the action (‘yes’ vs. ‘no’) in both hands, yet they do so in a different manner for each hand. Thus, the neural representation of semantic action goals - ‘why’ an action is performed - is encapsulated with the physical features of the action (‘how’ the action is performed). Alternatively, the regions of overlap use the same neural representation to differentiate ‘yes’/‘no’ answers for the two hands, but the lower SNR in left hand trials, as described above, did not allow us to detect such generalization across hands. This should be further tested and directly examined in future studies.

There is an ongoing debate as to whether conceptual information is stored in a-modal symbolic representations or grounded in sensory-motor simulations (Mahon and Caramazza 2008, Caramazza et al. 2014), as suggested by the embodied cognition hypothesis (Gallese and Lakoff, 2005). Previous neuroimaging studies supporting the embodied cognition hypothesis present evidence of neural representation of abstract action concepts (e.g. grasping, tearing, tossing), or manipulable artifact, which is shared across various physical exemplars (Barsalou, 2008; van Elk et al., 2014; Turella et al., 2020). However, to date there is no clear evidence as to whether introspective semantic goals, and not only the action, object or tool that is physically manipulated, are embodied and underlined by sensory-motor responses. Our results indicate that beyond the neural representation of performed actions and their sensory targets, semantic goals as well, are encoded within sensory-motor regions, in agreement with theories of grounded cognition (Gallese, 2003, 2009; Engel et al., 2013).

Localizing the neural circuits that underlie action goal representation has implications for the development of more accurate and efficient Brain-Machine Interfaces (BMIs) (Ortiz-Rosario and Adeli, 2013;Rezeika et al., 2018) such that genuine real-time human intentions (instead of memorized movement trajectories or visual cues) can be decoded and used to operate neuroprosthetics with increasing functionality. Moreover, shedding light on the neural architecture of action organization and identifying the processes by which internal states are constructed into actions can inspire the development of models for human action recognition. Finally, elucidating the neurophysiological link between actions and their underlying goals may provide insight with respect to pathologies such as apraxia in which action goal representation is compromised (Grafton & Hamilton, 2007).

## Acknowledgments

This research was supported by the Israel Science Foundation (grant No.2392/19 to R.M) The authors thank lab members for constructive comments and fruitful suggestions.

## References

Barsalou LW (2008) Grounded cognition. Annual Review of Psychology. https://doi.org/10.1146/annurev.psych.59.103006.093639

Beauchamp MS, Martin A (2007) Grounding object concepts in perception and action: Evidence from fMRI studies of tools. Cortex. https://doi.org/10.1016/S0010-9452(08)70470-2

Benjamini Y, Hochberg Y (1995) Controlling the False Discovery Rate: A Practical and Powerful Approach to Multiple Testing. J. R. Stat. Soc. Ser. B. 1995:p. 289--300. Journal of the Royal Statistical Society B.

Brouillet T, Heurley L, Martin S, Brouillet D (2010) The embodied cognition theory and the motor component of “yes” and “no” verbal responses. Acta Psychologica 134: 310–317. https://doi.org/10.1016/j.actpsy.2010.03.003

Caramazza A, Anzellotti S, Strnad L, Lingnau A (2014) Embodied cognition and mirror neurons: A critical assessment. Annual Review of Neuroscience 37: 1–15. https://doi.org/10.1146/annurev-neuro-071013-013950

Eisenberg M, Shmuelof L, Vaadia E, Zohary E (2011) The representation of visual and motor aspects of reaching movements in the human motor cortex. Journal of Neuroscience 31: 12377–12384. https://doi.org/10.1523/JNEUROSCI.0824-11.2011

van Elk M, van Schie H, Bekkering H (2014) Action semantics: A unifying conceptual framework for the selective use of multimodal and modality-specific object knowledge. Physics of Life Reviews 11: 220– 250. https://doi.org/10.1016/j.plrev.2013.11.005

Engel AK, Maye A, Kurthen M, König P (2013) Where’s the action? The pragmatic turn in cognitive science. Trends in Cognitive Sciences. https://doi.org/10.1016/j.tics.2013.03.006

Fernandino L, Humphries CJ, Seidenberg MS, Gross WL, Conant LL, Binder JR (2015) Predicting brain activation patterns associated with individual lexical concepts based on five sensory-motor attributes. Neuropsychologia 76: 17–26. https://doi.org/10.1016/j.neuropsychologia.2015.04.009

Gallese V (2003) A neuroscientific grasp of concepts: From control to representation. Philosophical Transactions of the Royal Society B: Biological Sciences 358: 1231–1240. https://doi.org/10.1098/rstb.2003.1315

Gallese V (2009) Motor abstraction: A neuroscientific account of how action goals and intentions are mapped and understood. Psychological Research 73: 486–498. https://doi.org/10.1007/s00426-009-0232-4

Gallese V, Lakoff G (2005) The brain’s concepts: The role of the sensory-motor system in conceptual knowledge. Cognitive Neuropsychology 22: 455–479. https://doi.org/10.1080/02643290442000310

Gallivan JP, Chapman CS, Wolpert DM, Flanagan JR (2018) Decision-making in sensorimotor control. Nature Reviews Neuroscience 19: 519–534. https://doi.org/10.1038/s41583-018-0045-9

Gallivan JP, Johnsrude IS, Randall Flanagan J (2016) Planning Ahead: Object-Directed Sequential Actions Decoded from Human Frontoparietal and Occipitotemporal Networks. Cerebral Cortex 26: 708–730. https://doi.org/10.1093/cercor/bhu302

Gallivan JP, Adam McLean D, Smith FW, Culham JC (2011) Decoding effector-dependent and effector-independent movement intentions from human parieto-frontal brain activity. Journal of Neuroscience 31: 17149–17168. https://doi.org/10.1523/JNEUROSCI.1058-11.2011

Gertz H, Lingnau A, Fiehler K (2017) Decoding movement goals from the fronto-parietal reach network. Frontiers in Human Neuroscience 11. https://doi.org/10.3389/fnhum.2017.00084

Gilron R, Rosenblatt J, Koyejo O, Poldrack RA, Mukamel R (2017) What’s in a pattern? Examining the type of signal multivariate analysis uncovers at the group level. NeuroImage 146: 113–120. https://doi.org/10.1016/j.neuroimage.2016.11.019

Hamilton AF d.C, Grafton ST (2008) Action outcomes are represented in human inferior frontoparietal cortex. Cerebral Cortex 18: 1160–1168. https://doi.org/10.1093/cercor/bhm150

Hamilton AFDC, Grafton ST (2006) Goal representation in human anterior intraparietal sulcus. Journal of Neuroscience 26: 1133–1137. https://doi.org/10.1523/JNEUROSCI.4551-05.2006

Hasson U, Hendler T, Bashat D Ben, Malach R (2001) Vase or face? A neural correlate of shape-selective grouping processes in the human brain. Journal of Cognitive Neuroscience. https://doi.org/10.1162/08989290152541412

Kiefer M, Pulvermüller F (2012) Conceptual representations in mind and brain: Theoretical developments, current evidence and future directions. Cortex 48: 805–825. https://doi.org/10.1016/j.cortex.2011.04.006

Krasovsky A, Gilron R, Yeshurun Y, Mukamel R (2014) Differentiating intended sensory outcome from underlying motor actions in the human brain. Journal of Neuroscience 34: 15446–15454. https://doi.org/10.1523/JNEUROSCI.5435-13.2014

de Lange FP, Spronk M, Willems RM, Toni I, Bekkering H (2008) Complementary Systems for Understanding Action Intentions. Current Biology 18: 454–457. https://doi.org/10.1016/j.cub.2008.02.057

Mahon BZ, Caramazza A (2008) A critical look at the embodied cognition hypothesis and a new proposal for grounding conceptual content. Journal of Physiology Paris 102: 59–70. https://doi.org/10.1016/j.jphysparis.2008.03.004

Miall RC, Wolpert DM (1996) Forward models for physiological motor control. Neural Networks. https://doi.org/10.1016/S0893-6080(96)00035-4

Mollo G, Pulvermüller F, Hauk O (2016) Movement priming of EEG/MEG brain responses for action-words characterizes the link between language and action. Cortex 74: 262–276. https://doi.org/10.1016/j.cortex.2015.10.021

Moritz X, Wurm F, Lingnau A (2015) Behavioral/Cognitive Decoding Actions at Different Levels of Abstraction. https://doi.org/10.1523/JNEUROSCI.0188-15

Orlov T, Makin TR, Zohary E (2010) Topographic Representation of the Human Body in the Occipitotemporal Cortex. Neuron 68: 586–600. https://doi.org/10.1016/j.neuron.2010.09.032

Ortiz-Rosario A, Adeli H (2013) Brain-computer interface technologies: From signal to action. Reviews in the Neurosciences 24: 537–552. https://doi.org/10.1515/revneuro-2013-0032

Pulvermüller F, Fadiga L (2010) Active perception: Sensorimotor circuits as a cortical basis for language. Nature Reviews Neuroscience. https://doi.org/10.1038/nrn2811

Rezeika A, Benda M, Stawicki P, Gembler F, Saboor A, Volosyak I (2018) Brain–computer interface spellers: A review. Brain Sciences 8. https://doi.org/10.3390/brainsci8040057

Rosenblatt JD, Benjamini Y, Gilron R, Mukamel R, Goeman JJ (2019) Better-than-chance classification for signal detection. Biostatistics: 1– 16. https://doi.org/10.1093/biostatistics/kxz035

Schaller F, Weiss S, Müller HM (2017) EEG beta-power changes reflect motor involvement in abstract action language processing. Brain and Language 168: 95–105. https://doi.org/10.1016/j.bandl.2017.01.010

Stelzer J, Chen Y, Turner R (2013) Statistical inference and multiple testing correction in classification-based multi-voxel pattern analysis (MVPA): Random permutations and cluster size control. NeuroImage 65: 69–82. https://doi.org/10.1016/j.neuroimage.2012.09.063

Sulzer JS, Chib VS, Hepp-Reymond MC, Kollias S, Gassert R (2011) BOLD correlations to force in precision grip: An event-related study. Proceedings of the Annual International Conference of the IEEE Engineering in Medicine and Biology Society, EMBS: 2342–2346. https://doi.org/10.1109/IEMBS.2011.6090655

Tian X, Poeppel D (2010) Mental imagery of speech and movement implicates the dynamics of internal forward models. Frontiers in Psychology. https://doi.org/10.3389/fpsyg.2010.00166

Turella L, Rumiati R, Lingnau A (2020) Hierarchical Action Encoding Within the Human Brain. Cerebral Cortex 30: 2924–2938. https://doi.org/10.1093/cercor/bhz284

Wentura D (2000) Dissociative Affective and Associative Priming Effects in the Lexical Decision Task: Yes Versus No Responses to Word Targets Reveal Evaluative Judgment Tendencies. Journal of Experimental Psychology: Learning Memory and Cognition 26: 456–469. https://doi.org/10.1037/0278-7393.26.2.456

Wolpert DM, Flanagan JR (2001a) Motor prediction. Current biology: CB. https://doi.org/10.1016/s0960-9822(01)00432-8

Wolpert DM, Flanagan JR (2001b) Primer Motor prediction. Current Biology 11: R729–32.

Wolpert DM, Ghahramani Z, Jordan MI (1995) Wolpert, D. M., Ghahramani, Z., & Jordan, M. I. (1995). An internal model for sensorimotor integration. Science-AAAS-Weekly Paper Edition, 269(5232), 1880–1882. Science-AAAS-Weekly Paper Edition.

Wurm MF, Ariani G, Greenlee MW, Lingnau A (2016) Decoding Concrete and Abstract Action Representations During Explicit and Implicit Conceptual Processing. Cerebral Cortex 26: 3390–3401. https://doi.org/10.1093/cercor/bhv169

Yarkoni T, Barch DM, Gray JR, Conturo TE, Braver TS (2009) BOLD correlates of trial-by-trial reaction time variability in gray and white matter: A multi-study fMRI analysis. PLoS ONE 4. https://doi.org/10.1371/journal.pone.0004257

